# A Hidden Markov Modeling Approach for Identifying Tumor Subclones in Next-Generation Sequencing Studies

**DOI:** 10.1101/675512

**Authors:** Hyoyoung Choo-Wosoba, Paul S. Albert, Bin Zhu

## Abstract

Allele-specific copy number alteration (ASCNA) analysis is for identifying copy number abnormalities in tumor cells. Unlike normal cells, tumor cells are heterogeneous as a combination of dominant and minor subclones with distinct copy number profiles. Estimating the clonal proportion and identifying mainclone and subclone genotypes across the genome is important for understanding tumor progression. Several ASCNA tools have recently been developed, but they have been limited to the identification of subclone regions, and not the genotype of subclones. In this paper, we propose subHMM, a hidden Markov model-based approach that estimates both subclone region as well as region-specific subclone genotype and clonal proportion. We specify a hidden state variable representing the conglomeration of clonal genotype and subclone status. We propose a two-step algorithm for parameter estimation, where in the first step, a standard hidden Markov model with this conglomerated state variable is fit. Then, in the second step, region-specific estimates of the clonal proportions are obtained by maximizing region-specific pseudo-likelihoods. We apply subHMM to study renal cell carcinoma datasets in The Cancer Genome Atlas. In addition, we conduct simulation studies that show the good performance of the proposed approach. The R package is available online at https://dceg.cancer.gov/tools/analysis/subhmm. somatic copy number alteration; tumor heterogeneity; E-M algorithm; forward-backward algorithm.

## 1 Introduction

Somatic copy number alterations (SCNA) are genetic changes in the cancer genome where the copy number of tumor cells departs from two copies through copy number deletion or amplification. SCNA analysis is used to identify genes with an abnormal copy number that contributes to carcinogenesis and cancer progression (Zack et al., 2013).

There are a number of analytical challenges in studying SCNA (Egeblad et al., 2010). First, the average copy number across chromosomes, called ploidy, may not be two and has to be estimated and adjusted for in the analysis. Second, there are various types of SCNA, some of which share the same total copy number. To distinguish them, allele-specific copy number alteration (ASCNA) analysis is preferred, which infers both the total copy number and the minor copy number in each locus. ASCNA analysis is essential in identifying copy-neutral LOH (loss of heterozygosity), in which one of the heterogeneous alleles is lost and the other is duplicated. (e.g., the genotype altering from AB to AA with the total copy number unchanged). Third, tumor tissue generally contains a mixture of tumor and normal cells; this type of heterogeneity, called tumor purity, needs to be accounted for to obtain correct inference about ASCNA. Finally, tumor cells themselves are heterogeneous with distinct copy number profiles reflecting a dominant clone (mainclone) and minor clones (subclone). Identifying main- and sub-clonal ASCNA across the genome would help design targeted therapeutics against genes with SCNA in both mainclone and subclone.

Several ASCNA tools, such as ASCAT (Van Loo et al., 2010), GPHMM (Li et al., 2011), MixHMM (Liu et al., 2010), and hsegHMM (Choo-Wosoba et al., 2018) have been developed that account for both tumor purity and ploidy. However, these methods assume that the tumor is homogeneous and consists of only a single clone. Tumor heterogeneity has been addressed by a number of authors ((Ha et al., 2014), (Shen and Seshan, 2016), (Li and Xie, 2015)), but no methods have been developed to infer multi-clonal genotype. We propose subHMM based on a hidden Markov model, that identifies multiple clones cross the genome. Standard HMM modeling has been used for both germline and somatic copy number variation (CNV) analysis ((Titsias et al., 2016), (Fan et al., 2017), (Cheng et al., 2017), (Yau et al., 2011), etc). In these cases, the forward-backward algorithm (Baum, 1972) with an E-M algorithm can be applied so that maximum-likelihood estimation is feasible. For the current problem where a clonal-specific genotype need to be identified across the genome, the forward-backward algorithm can not be used since the E-step involves computing conditional expectations of complex functions of sequential subclone indicators. Instead, a novel two-step algorithm is proposed for parameter estimation.

The remaining of the paper is organized as follows. In Section 2, we propose subHMM based on a hidden Markov model through the proposed two-step estimation procedure for inference. In Section 3, we apply subHMM to whole exome sequencing (WES) renal cell carcinoma data in the Cancer Genome Atlas (TCGA) project. We perform simulation studies to demonstrate the utility of the approach in Section 4. A discussion follows in Section 5.

## 2 Hidden Markov Model for Subclone: subHMM

Let *W_k_* be a hidden state of the *k*th locus representing mainclone genotype, subclone status, and subclone genotype, where *W_k_* follows a Markov chain with a transition probability *A*: *A_w′w_* = *P*(*W_k_* = *w*|*W*_*k*−1_ = *w′*) for *k* = 1, ⋯, *N* with the number of loci, *N*. More formally, we decompose *W_k_* into three different variables: *Z_k_*, *U_k_*, and *T_k_*. *Z_k_* indicates the mainclone genotype of the *k*th locus; *U_k_* is the indicator of whether subclone exists at the kth locus; *T_k_* is the subclone genotype when subclone exists. Figure 1 provides a schematic diagram detailing the transitions of *W_k_* across consecutive loci.

**Figure 1:**
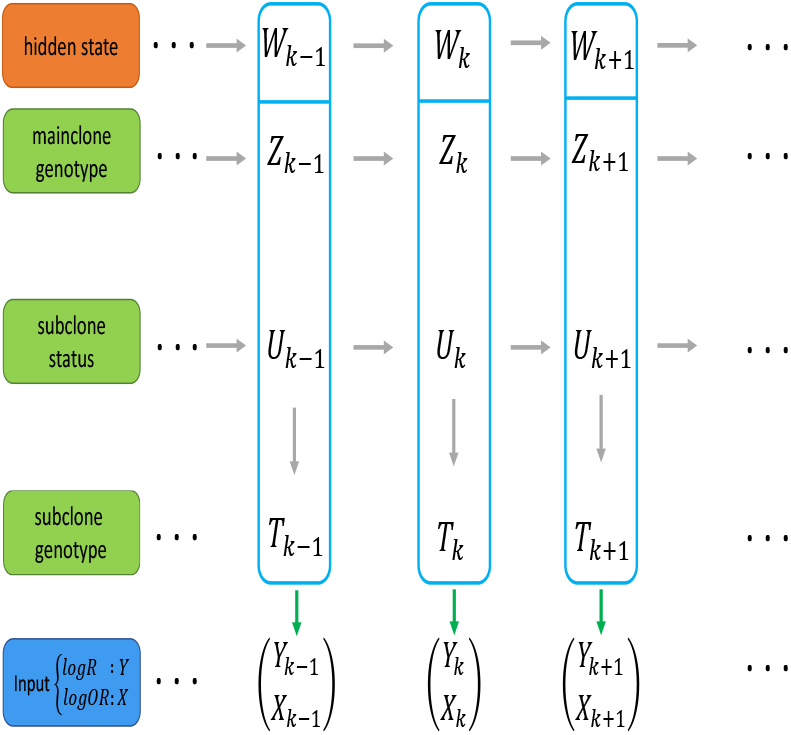
The transition flow chart of subHMM

We consider genotypes (described in Web Appendix A) up to total copy number five. The hidden state space indicator *W_k_* = {*Z_k_, U_k_, T_k_*} includes these twelve different mainclone genotypes, *Z_k_* ∈ G = {0, *A*, ⋯, *AAAAA*}, the indicator of a subclone *U_k_* ∈ {0, 1}, and eleven possible subclone genotypes, *T_k_* ∈ {*G*; *T_k_* ≠ *Z_k_*} when a subclone exists (*U_k_* = 1). For instance, *Z_k_* = *A*, *U_k_* = 1, and *T_k_* = *AAB* indicates that the mainclone genotype is *A* with the subclone genotype *AAB*; *Z_k_* = *AAA* and *U_k_* =0 means the mainclone genotype *AAA* with no subclone at the kth locus. Note that *T_k_* is not defined when *U_k_* = 0.

For the subclone genotype, we assume that the subclone genotype *T_k_* is different from *Z_k_*, the mainclone genotype, and *T_k_* follows a multinomial distribution. Then, the transition probability *A* can be expressed as

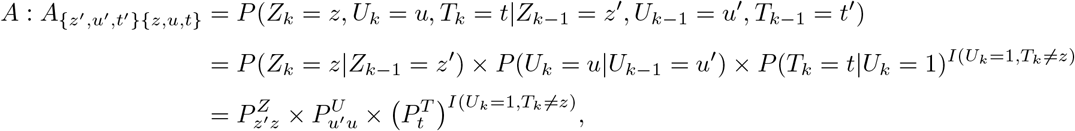

where 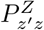 and 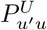 are elements in the probability transition matrices for *Z_k_* and *U_k_*, and 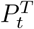 is the probability of *T_k_* = *t* where *t* ≠ *z*.

In practice, we do not observe the hidden states, rather we observe allele-specific measurements of logR and logOR, where logR characterizes the ratio of total copy numbers in tumor vs normal and logOR measures the odds ratio of copy numbers from maternal and paternal alleles in tumor vs normal. Consequently, these measurements depend on mainclone genotype as well as the status of subclones and their genotypes, detailed below.

Let *Y_k_* and *X_k_* be the *k*th observation of logR and logOR, respectively. Then, given the hidden states, we specify the conditional distributions of *Y_k_* and *X_k_*, separately. For *Y_k_*, we use a t-distribution with degrees of freedom to account for hypersegmentation that is common in next-generation sequencing (NGS)-based data ((Peel and McLachlan, 2000),(Choo-Wosoba et al., 2018)). Specifically, we use a normal-gamma mixture to derive the t-distribution with degrees of freedom (i.e. we assume *Y_k_* ~ *N*(*μ*, *κ*^2^/*δ*) with 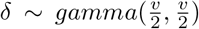 where *Y_k_* has a marginal expectation of *μ* with a variance of 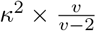). We assume that the squared logOR, 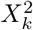, follows a non-central chi-square distribution with one degree of freedom and a non-central parameter *η* = *λ*^2^/*σ*^2^, where *λ* and *σ*^2^ are the mean and variance of logOR. Finally, the expectations of *Y_k_* and 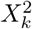 are defined as

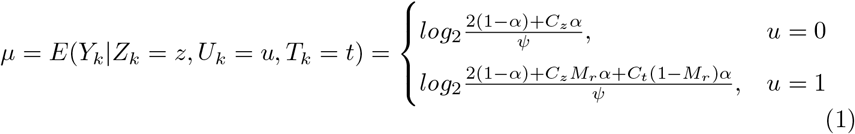

and

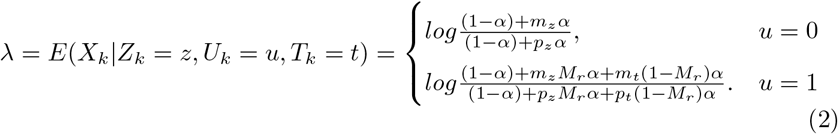

In Equations 2.1 and 2.2, *α* and *ψ* are the tumor purity and the average ploidy, respectively. A subclone region is delinerated along the chromosome where *U_k_* = 1 for consecutive loci. The parameter *M_r_* indicates the mainclone (or clonal) proportion of tumor cells in the rth subclone region. *C_z_* and *C_t_* are copy numbers from maternal and paternal alleles, *m_z_* and *m_t_* are copy numbers of maternal alleles, and *p_z_* and *p_t_* are copy numbers of paternal alleles, each corresponding to the mainclone (subscript *z*) and subclone (subscript *t*), respectively. *α*, *ψ*, *κ*^2^, *v*, and *σ*^2^ are global parameters over the entire genome.

Since logR and logOR values are assumed to be conditionally independent given the states, the joint emission probability is constructed simply by multiplying the two density functions.

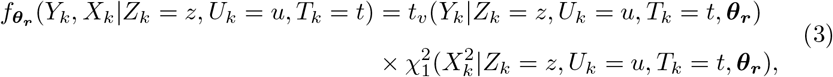

where ***θ_r_*** = {*α*, *ψ*, *M_r_*, *κ*^2^, *σ*^2^, *v*}.

Based on this model structure, we consider two different assumptions about the clonal proportion. First, we assume a constant clonal proportion where all the identified sublone regions have the exact same clonal proportions of tumor cells. Second, we assume that each subclone region has a potentially different clonal proportion (*M_r_*). The first model is consistent with a single subclone, and the second with multiple subclones in the tumor.

### 2.1 subHMM with E-M algorithm under the constant clonal proportion assumption

We consider the situation where mainclone/subclone proportions are constant across all the subclone regions (*M_r_* = *M*). Since *W_k_* follows an unobserved Markov chain, we can directly apply the E-M algorithm to estimate all the parameters by maximizing the expectation of a complete log likelihood function

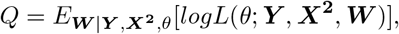

where ***θ_r_*** = {*α*, *ψ*, *M*, *κ*^2^, *σ*^2^, *v*}, and where we denote all the bold characters as vectors, such as ***Y*** = {*Y*_1_, *Y*_2_, ⋯, *Y_N_*}, ***X***^2^ = {*X*_1_, *X*_2_, ⋯, *X_N_*}, and *W* = {*W*_1_, *W*_2_, ⋯, *W_N_*}. Rewriting ***W*** with respect to ***Z, U***, and ***T***, our expectation of the complete log likelihood function *Q* is then redefined as,

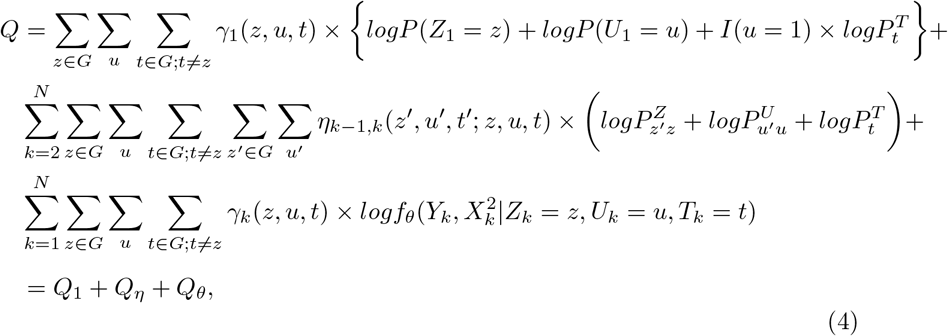

where *γ_k_*(*z, u, t*) = *P*(*Z_k_* = *z*, *U_k_* = *u, T_k_* = *t*|***Y, X***^2^) and *η*_*k*−1,*k*_(*z′*, *u′*, *t′*; *z, u, t*) = *P*(*Z_k_* = *z*, *U_k_* = *u*, *T_k_* = *t*, *Z*_*k*−1_ = *z′*, *U*_*k*−1_ = *u*′, *T*_*k*−1_ = *t′*|***Y, X***^2^) and importantly, the indicators are sufficient statistics for initial probabilities, transition probabilities, and emission probability.

In the E-step, we obtain *γ_k_* (*z, u, t*) and *η*_*k*−1,*k*_ (*z′, u′, t′; z, u, t*) with the forward-backward algorithm introduced by Baum (1972). The forward algorithm starts with calculating the forward element of the first loci measurements, *a_w_*(1), de-composed to *Z*, *U*, and *T*, following by

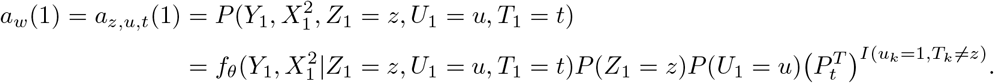

Based on *a_z,u,t_*(1), the forward algorithm obtains sequential elements corresponding to each of *k* locus given by

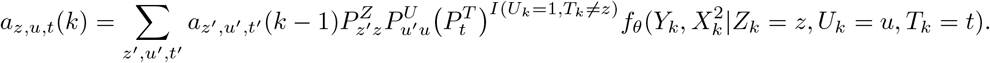

The backward element of the *k*th locus is defined as

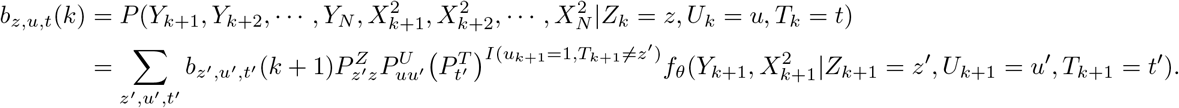

The backward algorithm starts with calculating the backward element of the last locus, *b_z,u,t_*(*N*) defined as 1 for all the hidden states, and obtains all the backward elements recursively. Finally, we can compute *γ_k_*(*z,u,t*) and *η*_*k*−1,*k*_(*z′, u′, t′*; *z,u,t*) by using both forward and backward elements, given by

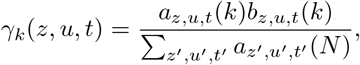

and

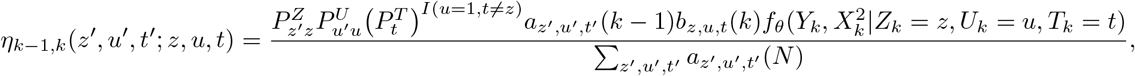

which are both sufficient statistics of *Q* (Equation 2.4) under the constant *M* assumption.

In E-step, given *γ_k_*(*z, u, t*) and *η*_*k*−1,*k*_(*z*_1_, *u*_1_, *t*_1_; *z*, *u, t*), we can finally estimate all the parameters by maximizing *Q*. First, we estimate 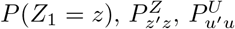, and 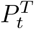 by closed forms. For *P*(*Z*_1_ = *z*),

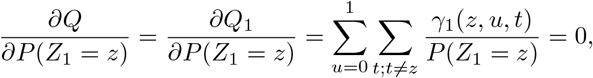

with respect to 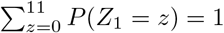. According to Lagrangian multipliers,

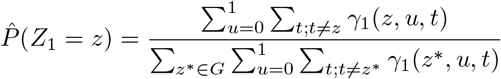

Similarly, for *P*(*U*_1_ = *u*),

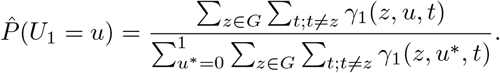

For estimating 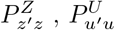, and 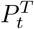,

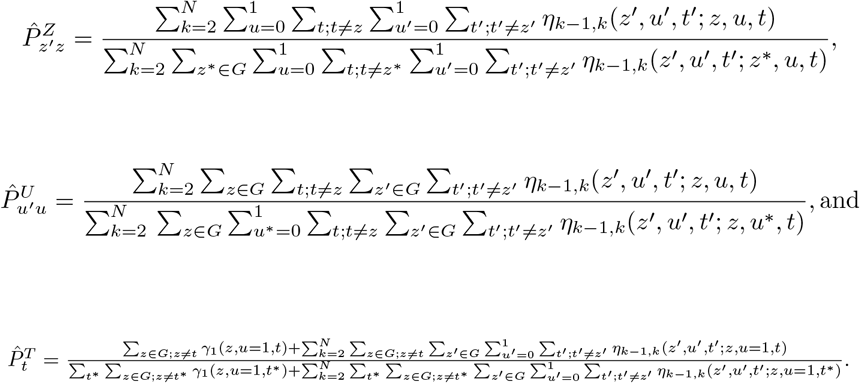

We iterate this E-M procedure until convergence to obtain the final global estimates. Specifically, all the global parameters including *M* are estimated by using *optim* function in R for the M-step and the forward-backward algorithm for the E-step. Mainclone genotypes are identified by choosing the maximum posterior probability at each locus. Subclone regions are identified by consecutive loci for which a state with a subclone (*U_k_* = 1) obtains the maximum posterior probability. To estimate the region-specific subclone genotype distribution, for each subclone region, we average *γ_k_*(*z, u* = 1, *t*), the posterior probabilities of each subclone genotype over all the possible mainclone genotypes with the subclone genotype. We determine the subclone genotype at the *r*th region by choosing the genotype with the largest average posterior probability.

### 2.2 subHMM with a two-step algorithm under the different clonal proportion assumption

subHMM that incorporates a varying clonal proportion could be formulated with an emission distribution, *f_θ_* (the third line in Equation 2.4), that depends on a region-specific clonal proportion. However, the complete data likelihood would be a complex function of the sequence of subclonal latent variables, *U_k_*. Because of this, a forward-backward algorithm cannot be implemented, making E-step computations in the E-M algorithm intractable. Hence, we propose a two-step algorithm that accounts for different clonal proportion across subclonal regions.

#### 2.2.1 The first step

We first assume that the clonal proportion is constant across subclone regions, *M_r_* = *M*. Under this constant clonal proportion assumption, we estimate the global parameters, the probability transition matrix corresponding to the mainclone, and identify the subclone regions as described in Section 2.1.

#### 2.2.2 The second step

For each subclone region (identified in the first step), we estimate region-specific clonal proportions and subclone genotype probability distribution. The clonal proportion of the *r*th subclone region, 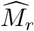 is estimated by locally maximizing the conditional expectation of the log likelihood function for the *r*th region *R_r_*, given all the global parameters estimated at the first step:

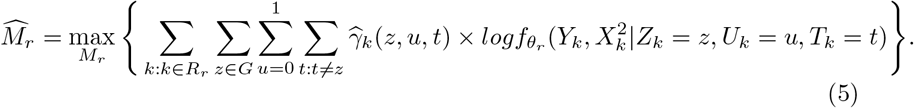

### 2.3 Asymptotic standard errors of global estimators

The Hessian-based standard errors require calculating the full log likelihood, which is only obtainable under the constant clonal proportion assumption. For the case of different clonal proportions, we can only compute the log likelihood in the first step of the two-step algorithm under the constant clonal assumption. Therefore, it is important to justify the use of Hessian-based standard errors under the assumption of different clonal proportions in subclone regions (see details in Section 2.2).

Under the constant clonal assumption, the log likelihood function given the estimates is then able to be obtained from the forward-backward algorithm (Section 2.1) by 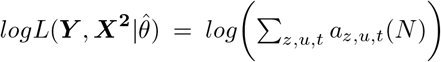, where *a_z,u,t_*(*N*) is the forward function of the last data point at each hidden state. Then, the asymptotic standard errors of the global estimators, based on the Hessian matrix is calculated as

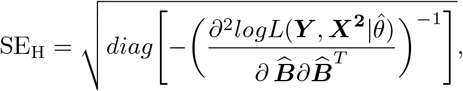

where ***B*** = (*α, ψ, κ*^2^, *v, σ*^2^)*^T^* is the vector of global parameters, and the Hessian-based standard error SE_H_ can be implemented with Hessian function in the R package *numDeriv*.

## 3 Analysis of Renal Cell Carcinoma Data

Initially, we apply subHMM to a single renal cell carcinoma sample with ID, TCGA-KL-8331 from TCGA project (https://cancergenome.nih.gov). TCGA project holds abundant cancer molecular profiling information to understand cancer genomics from a molecular perspective, including large-scale genome sequencing data. The whole exome sequencing dataset of TCGA-KL-8331 is available at https://portal.gdc.cancer.gov, that contains read counts and read depths over entire chromosomes for both normal and tumor samples within the same patient. We use preprocessing codes in FACETS (Shen and Seshan, 2016) (preProcSample, and procSample) to obtain logR and logOR measurements from the read-based dataset. A thinning procedure is then conducted by keeping every 10th values of logR and logOR to reduce excessive numbers of copy number changes caused by sequencing artefacts (hypersegmentation) and to reduce the computation burden, resulting in 36,914 pairs of logR and logOR. Recalling the definition of logOR, if a locus is homozygous, the odds ratio is 0 and logOR is undefined and only logR measurements contribute the estimation of ASCNA. For the sample TCGA-KL-8331, 4,691 (approximately 13% of 36,914 pairs) loci are heterozygous across the whole chromosomes, for which logOR values are defined.

Figure 2 shows the copy number profile of all the chromosomes based on subHMM with the hidden state space reflecting copy numbers of up to 5. Figure 2.a presents logR (the top plot) and logOR (the bottom plot) values across the genome with blue dots. The red lines indicate the estimated segment mean of logR and logOR based on subHMM. Figure 2.b shows estimated clonal profiles of mainclone genotypes (the top plot), subclone regions (the middle plot), and subclone genotypes (the bottom plot). The most frequent identified mainclone genotypes are A (deletion event) and AB (normal status) across the genome. Figure 2.b shows 5 subclone regions identified at chromosomes 11, 18, 22, and chromosome X. The subclone regions are also shown as green dots in Figure 2.a. The subclone genotype distributions for regions 1 to 4 suggest that the subclone genotypes are equally likely to be AB or AABB; No genotype is clearly favored for region 5 (Table B.1 in Appendix B of the Supplementary Materials).

**Figure 2:**
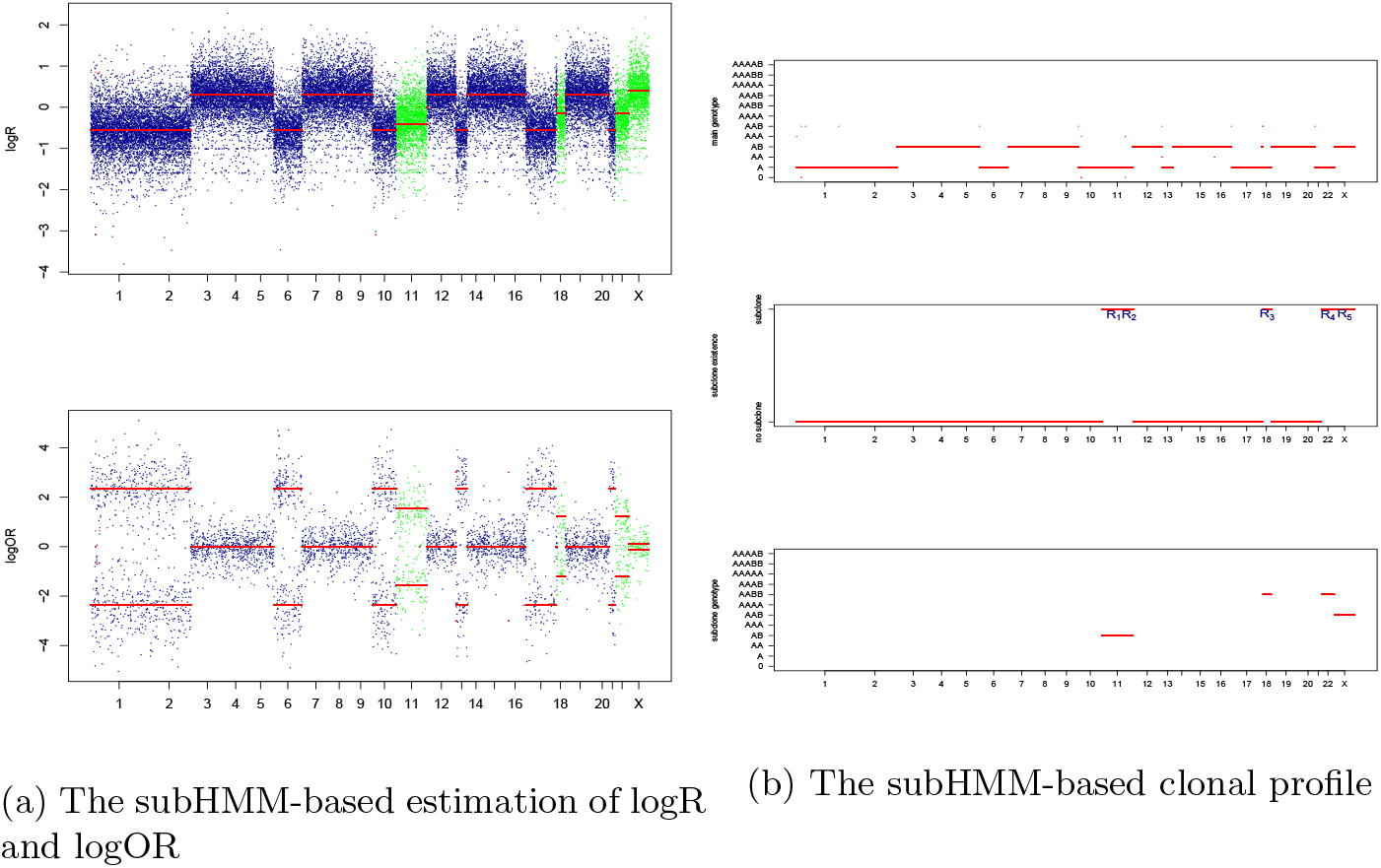
subHMM-based subclonal profile of TCGA-KL-8331; the rth subclone region *R_r_* defines as a consecutive loci where *U_k_* = 1 with at least a length of 50

Table 1 summarizes the result of both the global parameter and clonal proportion parameter estimation. Specifically, the estimated purity 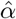 is 0.91, which indicates that the tumor sample contains 91% tumor cells. The estimated ploidy below 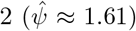 refers to partial loss of chromosomes in the tumor sample. The clonal proportions obtained by Equation 2.5 for each of the identified subclone regions, *M*_1_ to *M*_5_ are also presented in Table 1. Since the region-specific clonal proportions are estimated by maximizing a localized likelihood, Hessian-based asymptotic standard error cannot easily be obtained. Instead, we propose a parametric bootstrap for the estimation of these standard errors. First, we simulate a sample under the model using the estimates as known parameters. Then, we fit subHMM to the simulated data. Standard errors for the estimated clonal proportions (SE_bs_ in Table 1) are obtained by repeating the bootstrap procedure 300 times, and obtaining the empirical standard deviations of the clonal proportion estimates. All the five estimated clonal proportions including the constant proportion *M* (obtained in the first step) range from 84% to 89%, which suggests that the proportion of subclone cells in tumor cells ranges between 11% to 16%.

**Table 1:**
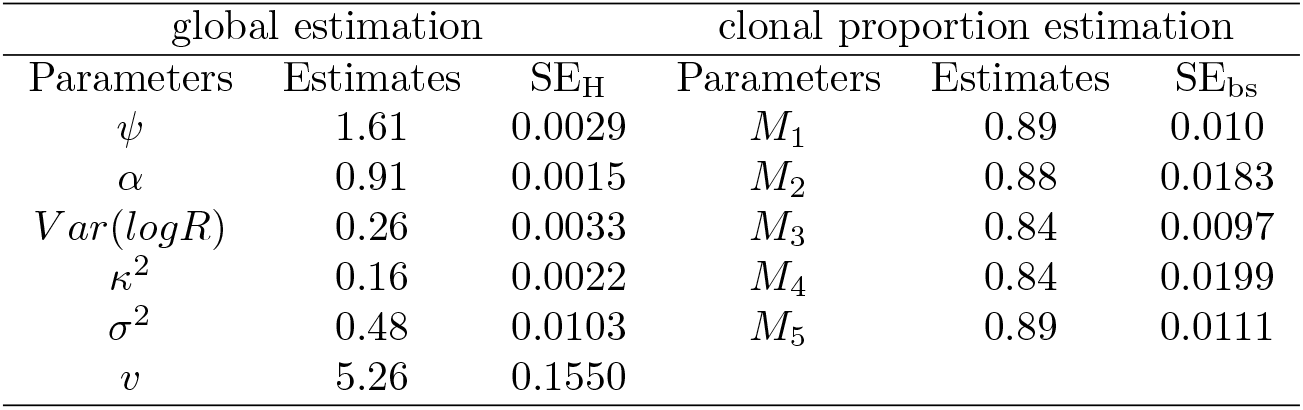
Global parameter estimation results of TCGA-KL-8331 dataset with subHMM based on state space of copy number 5

| global estimation |  |  | clonal proportion estimation |  |  |
| --- | --- | --- | --- | --- | --- |
| Parameters | Estimates | SE<sub>H</sub> | Parameters | Estimates | SE<sub>bs</sub> |
| $\psi$ | 1.61 | 0.0029 | $M_1$ | 0.89 | 0.010 |
| $\alpha$ | 0.91 | 0.0015 | $M_2$ | 0.88 | 0.0183 |
| $Var(logR)$ | 0.26 | 0.0033 | $M_3$ | 0.84 | 0.0097 |
| $\kappa^2$ | 0.16 | 0.0022 | $M_4$ | 0.84 | 0.0199 |
| $\sigma^2$ | 0.48 | 0.0103 | $M_5$ | 0.89 | 0.0111 |
| $v$ | 5.26 | 0.1550 | | | |

As shown in Figure 2.b, the predominant (main) genotypes are A and AB, suggesting that the state-space of the HMM may be reduced to (0, A, AA, AB) for both the mainclone and subclone genotypes, corresponding to a maximum copy number of two. Penalized likelihood techniques such as AIC and BIC (shown Table B.3 in Appendix B of the Supplementary Materials) show that a reduced model with a maximum copy number of 2 for both the main-and sub-clone genotypes is favored over models with a maximum copy number of 5. Figure 3 shows four subclone regions identified with this reduced model. These regions are similar to those identified with the complex model with the exception of an additional region identified on chromosome 5 and the lack of a region on chromosome X. The region-specific subclone genotypes presented in the Appendix B of the Supplementary Materials (Table B.2) suggests that the regions *R*_2_ to *R*_4_ have an AB genotype with high confidence and the region *R*_1_ has a genotype of A or AA with equal confidence.

**Figure 3:**
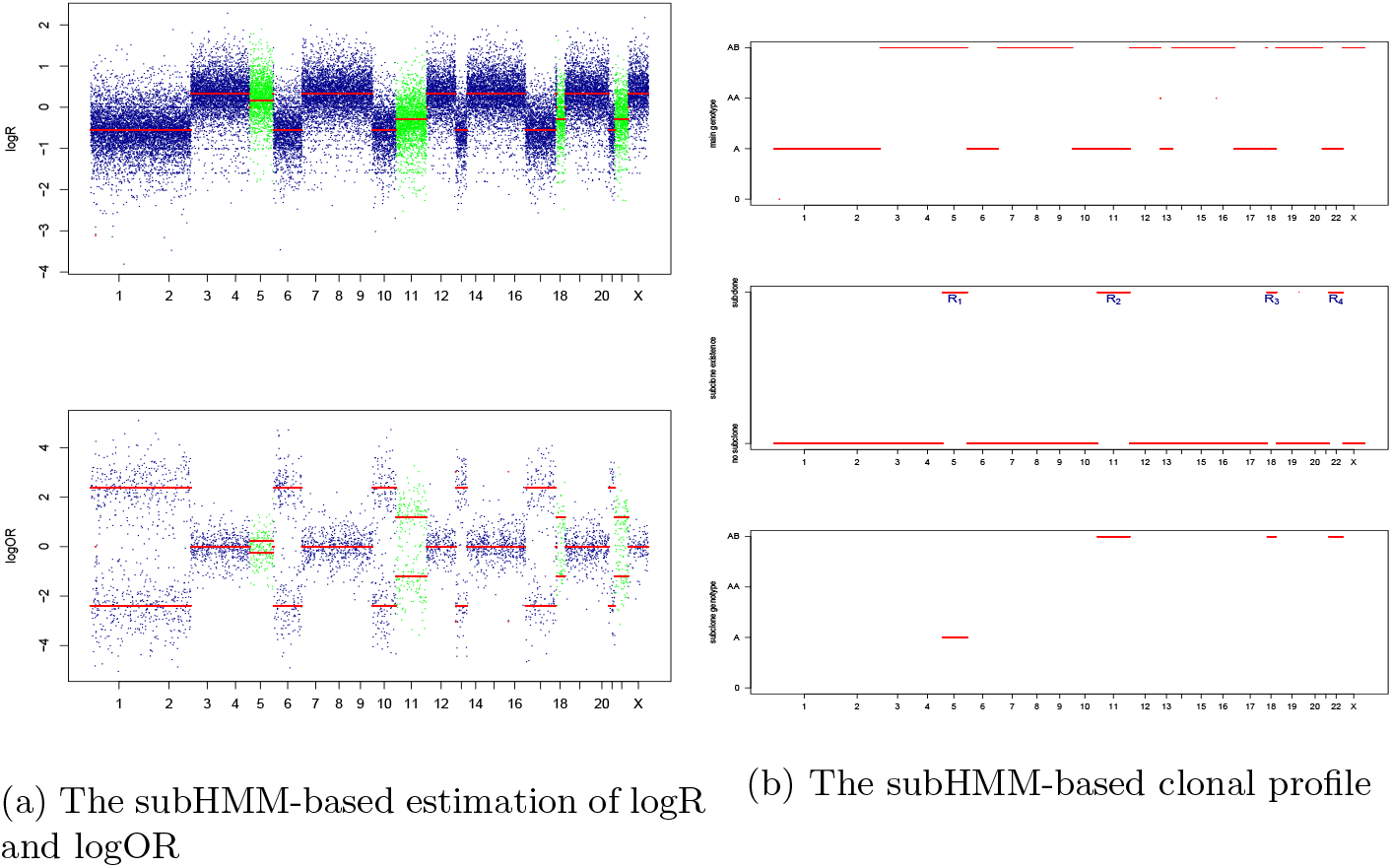
subHMM-based subclonal profile of TCGA-KL-8331 with copy number upto 2 as hidden state space; the rth subclone region *R_r_* defines as a consecutive loci where *U_k_* = 1 with at least a length of 50

Estimated global parameters for the reduced model are presented in Table 2. These estimates are similar to those estimated for the more complex model (Table 1). In contrast with the global parameters, the region-specific clonal proportions (presented in Table 2) show more variation between regions than those estimated with the more complex model. The clonal proportion estimates *M*_3_ and *M*_4_, have larger standard errors than those estimated for *M*_1_ and *M*_2_, possibly due to their regions being shorter in length. The estimates of *M*_1_, *M*_2_, *M*_3_, and *M_4_* along with their associated standard errors (obtained with the bootstrap) suggest that there may be two underlying clonal proportions; one at 0.7 and the other 0.8. These two distinct values suggest that there are two subclone events in this TCGA-KL-8331 sample. Further evidence of at least two subclones is obtained with the use of a bootstrap Wald test (null hypothesis of constant clonal proportion) that was highly significant (*p* < 0.0001). This new insight on region-specific clonal proportions and genotype is not available with other approaches ((Shen and Seshan, 2016): Appendix C of the Supplementary Materials shows that the subclone regions and mainclone genotypes are similar to those obtained with our approach).

**Table 2:**
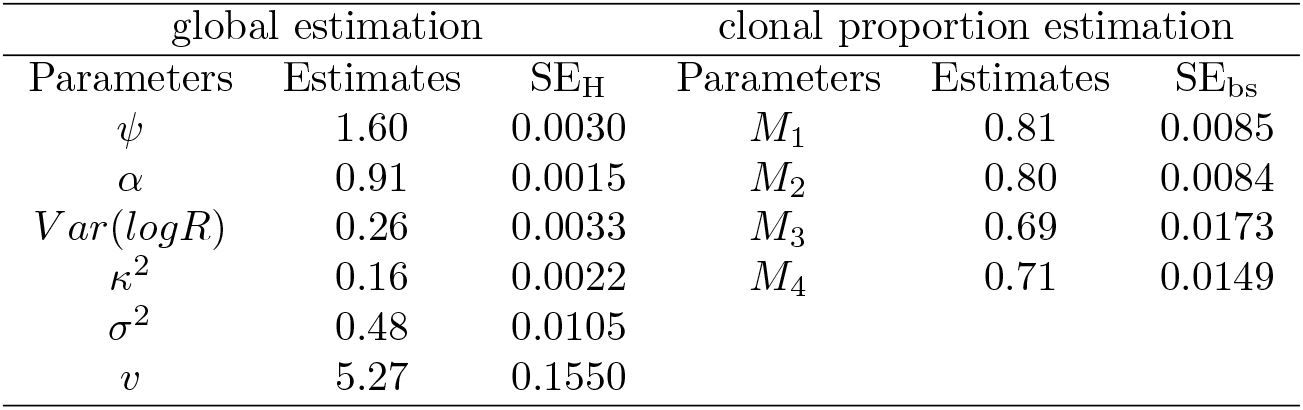
Global estimation results of TCGA-KL-8331 dataset with subHMM based on state space of copy number 2

In practice, it is of interest to apply subHMM to a number of samples in order to characterize recurrent mainclone or subclone SCNA events across samples. We apply subHMM to 316 renal cell carcinoma samples from TCGA. Using AIC, we compared subHMM with a model hsegHMM that assumes the existence of mainclone only in tumor sample (Choo-Wosoba et al., 2018); 120 of 316 showed a lower AIC for subHMM compared to hsegHMM, suggesting the presence of subclones. We focus on these 120 samples in the following analyses. We summarize the results of these subHMM-favored samples by creating a cytoband-based stacked histogram of allele-specific SCNA events. These histograms are used to examine the frequency of allele-specific SCNA events over samples, across prespecified chromosomal locations. We found that the most frequent mainclone SCNA event is a hemizygous deletion (genotype A) on Chromosomes 3 (Figure D.1 in Appendix D of the Supplementary Materials) and a copy number gain (genotype AAB) between cytoband q21.3 and q35.3 on Chromosome 5 (Figure D.1 in Appendix D of the Supplementary Materials). Figure D.2 in Appendix D of the Supplementary Materials shows subclone SCNA events across the entire chromosomes; the most frequent subclone SCNA event is whole-genome doubling with genotype AABB (balanced copy number amplification; 15 cases out of 120 are identified as the dominant subclone genotype AABB across the entire genome) as shown in Figures D.2 and D.3 in Appendix D of the Supplementary Materials. Identifying subclone SCNA events is a unique feature of our approach and cannot be done with previously proposed methods. The high rate of balanced copy number amplification among subclonal cells may be a defining feature of the tumor type and may suggest a new target therapy.

## 4 Simulations

We evaluate the performance of subHMM under the different clonal proportion scenario. Specifically, we investigate if the estimation of global parameters obtained at the first step is robust to an incorrect constant clonal proportion assumption that is made at this step in the two-step algorithm. We also evaluate the estimation of region-specific clonal proportions as well as subclone regions and their corresponding subclone genotypes.

All the results and summaries are based on 300 simulated datasets. Each dataset includes four subclone regions with clonal proportions (*M*_1_ = *M*_2_ = 0.8, *M*_3_ = *M*_4_ = 0.7) and has a similar copy number profile to the result from Section 3 (the application study) based on the state space corresponding to copy number 2. First, we specifiy three different probability matrices, 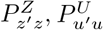, and 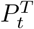 to form the probability transition matrix A. Second, we generate a sequence from a Markov chain with transition matrix *A* of size 30,000 by using the R package *markovchain*. Given this sequences, both the expectations of logR and logOR are computed by using Equations 2.1 and 2.2 with the given parameters. Finally, logR and logOR observations are randomly generated from t- and normal distributions with means given by their expected values and measured error variances, respectively. Similar to the application, 90% of logOR values are assigned as missing values. All the results and summaries are based on 300 simulated datasets.

According to Table 3, all the global estimates and corresponding Hessian-based standard errors (SE_H_) are close to the true values. This demonstrates that global parameter estimates and their associated Hessian-based standard errors obtained in the first step are unbiased even when the clonal proportions vary across regions. The clonal proportion estimation obtained from the second step is also unbiased (Table 4).

**Table 3:**
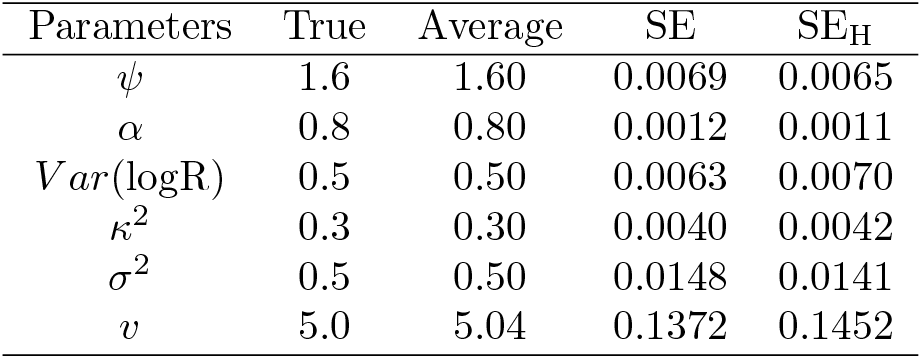
Global estimation result of the different clonal proportion assumption-based simulation study with subHMM based on 300 datasets

**Table 4:** Mainclone proportion simulation result of the different clonal proportion assumption-based simulation study with subHMM based on 300 datasets

| Parameters | True | Average | SD<sup>1</sup> |
| --- | --- | --- | --- |
| $M_1$ | 0.8 | 0.80 | 0.0063 |
| $M_2$ | 0.8 | 0.80 | 0.0076 |
| $M_3$ | 0.7 | 0.70 | 0.0102 |
| $M_4$ | 0.7 | 0.70 | 0.0097 |

<sup>1</sup> Monte-Carlo-based standard deviations

Furthermore, we examine the accuracy of genotypes. Figure 4 shows the probability of mainclone genotype, subclone region, and subclone genotype detection across the genome (black lines). The true clonal regions and genotypes are also shown (red lines). The figure demonstrates that subHMM does a good job in estimating these genomic features in realistic scenarios such as the renal cell carcinoma sample.

**Figure 4:**
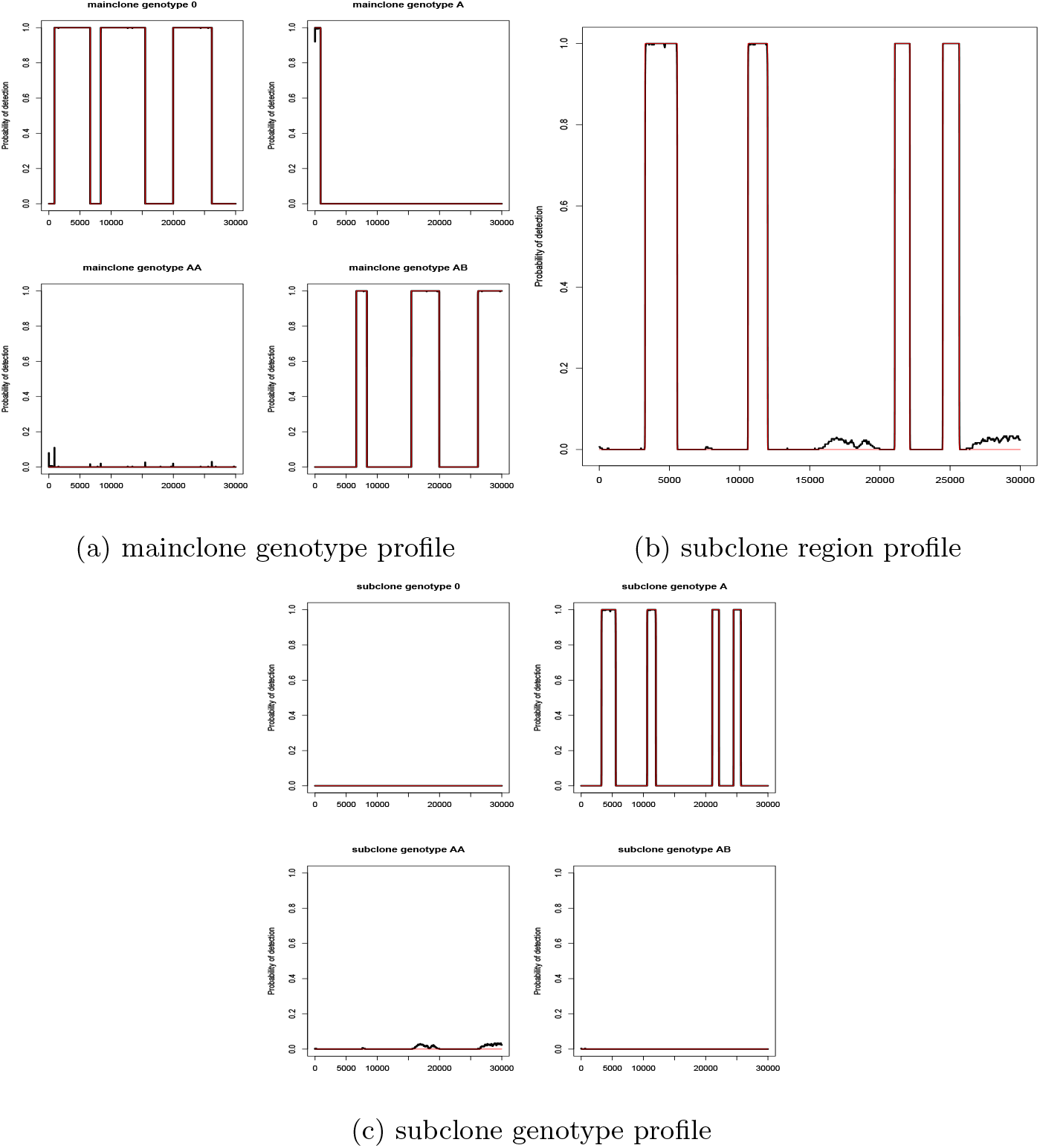
The probability of correct identification under the different clonal proportion assumption simulation study; The results are based on 300 simulated datasets; The red lines and black lines represent true and estimated profiles, respectively.

## 5 Discussion

In this paper, we developed a new hidden Markov model approach analyzing ASCNA that allows identifications of subclonal copy number alterations in heterogeneous tumor samples. Specifically, we use a hidden Markov model structure to locate these clonal-specific copy number alterations and their associated genotypes. Through this additional inference, our method provides further understanding of the carcinogenesis process that may ultimately lead to new cancer treatments by targeting subclonal copy number alterations of cancer driver genes.

We proposed a two-step estimation approach where we made the constant clonal proportion assumption in the first step and then estimate different proportions corresponding to identified subclone regions in the second step. Inference on the regional subclone proportions and their associated genotype is a challenging statistical problem. Specifically, when the clonal proportion varies by subclone, the emission distribution depends on the region-specific clonal proportion. In this case, the forward-backward algorithm cannot be applied, making E-step computations intractable. Inspired by this challenge, the two-step algorithm is proposed to handle this complex from intractable to tractable solution. In the first step, the standard forward-backward algorithm in HMM can evaluate the full log likelihood based on two sets of expected sufficient statistics, *γ_k_* and *η*_*k*−1,*k*_, under the constant clonal proportion assumption. The proposed methodology is simple to implement and will provide practitioners the opportunity for enhanced interence in SCNA.

As illustrated in this article, our methodology can be used for a population-based study with many samples. In our analysis, we summarized molecular profiles across samples of the same tumor. Other applications are possible, where we compare these SCNAs across tumor type, different types of exposure, or by different treatments.

## 6 Software

Software in the form of R code, together with a sample input data set and complete documentation is available as a R package at https://dceg.cancer.gov/tools/analysis/subhmm.

## 7 Supplementary Material

Supplementary material is available in a separate file.

## Acknowledgments

We would like to acknowledge our usage of the data from The Cancer Genome Atlas (TCGA) supported by the National Cancer Institute and National Human Genome Research Institute: https://cancergenome.nih.gov. We would like to thank Bill Wheeler (Information Management Services) and Lei Song (Biostatistics Branch) for computational contributions. *Conflict of Interest:* None declared.

