## Supplemental Tables and Figures for "A Hidden Markov Modeling Approach for Identifying Tumor Subclones in Next-Generation Sequencing Studies"

#### Web Appendix A. Description of tumor main clonal genotype states and corresponding genotype of total copy number

Table A.1: deletion (DEL), hemizygous deletion LOH (DLOH), copy neutral LOH (NLOH), diploid heterozygous (HET), gain of 1 allele (GAIN), amplified LOH (ALOH), allele-specific copy number amplification (ASCNA), balanced copy number amplification (BCNA), and unbalanced copy number amplification (UBCNA).

| State($z$) | Genotype | Copy Number ($C_z$) | Allelic information |
|---|---|---|---|
| 0 | 0 | 0 | HOMD |
| 1 | A | 1 | DLOH |
| 2 | AA | 2 | NLOH |
| 3 | AB | 2 | HET |
| 4 | AAB | 3 | GAIN |
| 5 | AAA | 3 | ALOH |
| 6 | AAAB | 4 | ASCNA |
| 7 | AABB | 4 | BCNA |
| 8 | AAAA | 4 | ALOH |
| 9 | AAAAB | 5 | ASCNA |
| 10 | AAABB | 5 | UBCNA |
| 11 | AAAAA | 5 | ALOH |

### Web Appendix B. Additional results of TCGA-KL-8331 dataset

Table B.1: The probabilities of subclone genotypes in identified subclone regions based on the state space of copy number 5

| regions | 0 | A | AA | AB | AAA | AAB | AAAA | AABB | AAAB | AAAAA | AAABB | AAAAB |
|---|---|---|---|---|---|---|---|---|---|---|---|---|
| $R_1$ | 0.01 | 0.00 | 0.00 | 0.50 | 0.00 | 0.04 | 0.00 | 0.43 | 0.00 | 0.00 | 0.01 | 0.00 |
| $R_2$ | 0.01 | 0.00 | 0.00 | 0.49 | 0.00 | 0.04 | 0.00 | 0.44 | 0.00 | 0.00 | 0.01 | 0.00 |
| $R_3$ | 0.01 | 0.00 | 0.00 | 0.46 | 0.00 | 0.04 | 0.00 | 0.46 | 0.00 | 0.00 | 0.01 | 0.00 |
| $R_4$ | 0.01 | 0.00 | 0.00 | 0.46 | 0.00 | 0.04 | 0.00 | 0.47 | 0.00 | 0.00 | 0.01 | 0.00 |
| $R_5$ | 0.03 | 0.14 | 0.21 | 0.00 | 0.11 | 0.27 | 0.02 | 0.11 | 0.07 | 0.00 | 0.02 | 0.01 |

Table B.2: The probabilities of subclone genotypes in identified subclone regions based on the state space of copy number 2

| regions | 0 | A | AA | AB |
|---|---|---|---|---|
| $R_1$ | 0.02 | 0.53 | 0.45 | 0.00 |
| $R_2$ | 0.04 | 0.00 | 0.03 | 0.93 |
| $R_3$ | 0.03 | 0.01 | 0.03 | 0.93 |
| $R_4$ | 0.03 | 0.00 | 0.03 | 0.94 |

Table B.3: Model selection criteria result of subHMM with three different state space cases

| | CN5/CN5[1] | CN2/CN2[2] | CN2/CN5[3] |
|---|---|---|---|
| $logL$ | -30680.99 | -30679.36 | -30667.15 |
| $AIC$ | 61905.99 | 61422.72 | 61462.31 |
| $BIC$ | 64222.43 | 61695.24 | 62007.36 |

[1] the complex model with the maximum copy number of five for both main-and sub-clone genotypes

[2] the complex model with the maximum copy number of two for both main-and sub-clone genotypes

[3] the complex model with the maximum copy number of two for main- and five for sub-clone genotypes

#### Web Appendix C. An ASCNA result with FACETS as an example of applying a current method

Figure C.1 shows the result of FACETS analysis on TCGA-KL-8331 dataset. The top panel represents the logR profile including the estimates (red lines), diploid state (purple line), and median (bright green line). The middle shows the logOR profile with the estimates (red lines) and the bottom presents the copy number profile with two different colored lines. The black lines indicate the total copy number and the red ones are the minor copy number. For example, the chromosomes 3 to 5 show the total copy number of two with one minor copy number, which refers to genotype AB.

It is clear that unlike subHMM capable of detecting region-specific subclone genotypes, FACETS only provides the identification of subclone regions (chr 11, 18 and 22) and the clonal proportions corresponding to those identified regions shown in the bottom pannel. Interestingly, our method identifies one more subclone region besides them, which indicates chr 5. The estimated hidden status in chr 5 is turned out to be the mainclone genotype AB with subclone region. However, the corresponding posterior probabilities are not dominantly larger than the second largest posterior probabilities whose corresponding hidden states indicate the same genotype of mainclone but with no subclone.

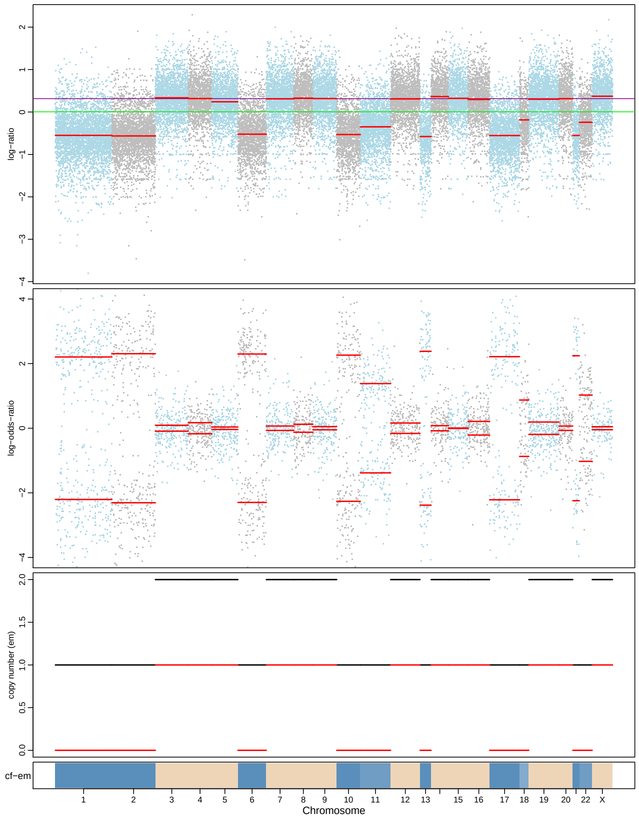


Figure C.1: FACETS results for the renal cell carcinoma sample, TCGA-KL-8331, after a thinning process.

### Web Appendix D. Population-based study

We perform ASCNA analysis of 120 samples out of 316 TCGA renal cell carcinoma samples, which are favored to hsegHMM (mainclone only assumption model, Choo-Wosoba et al., 2018) based on AIC. We summarize the results of the 120 samples by creating stacked histograms for the mainclone, subclone region, and subclone genotype profiles, respectively. The x-axis units refer to predefined cytobands corresponding to the genetic locations that subHMM identifies as ASCNA.

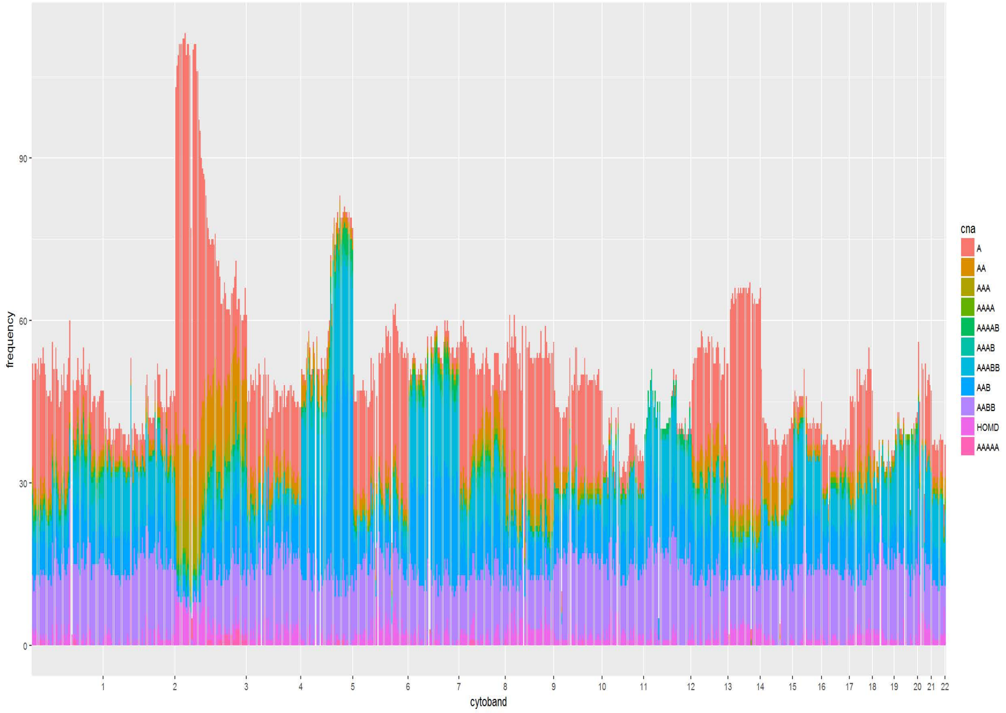


Figure D.1: Stacked histogram of mainclone genotype profile based on subHMM for 120 TCGA renal cell carcinoma samples

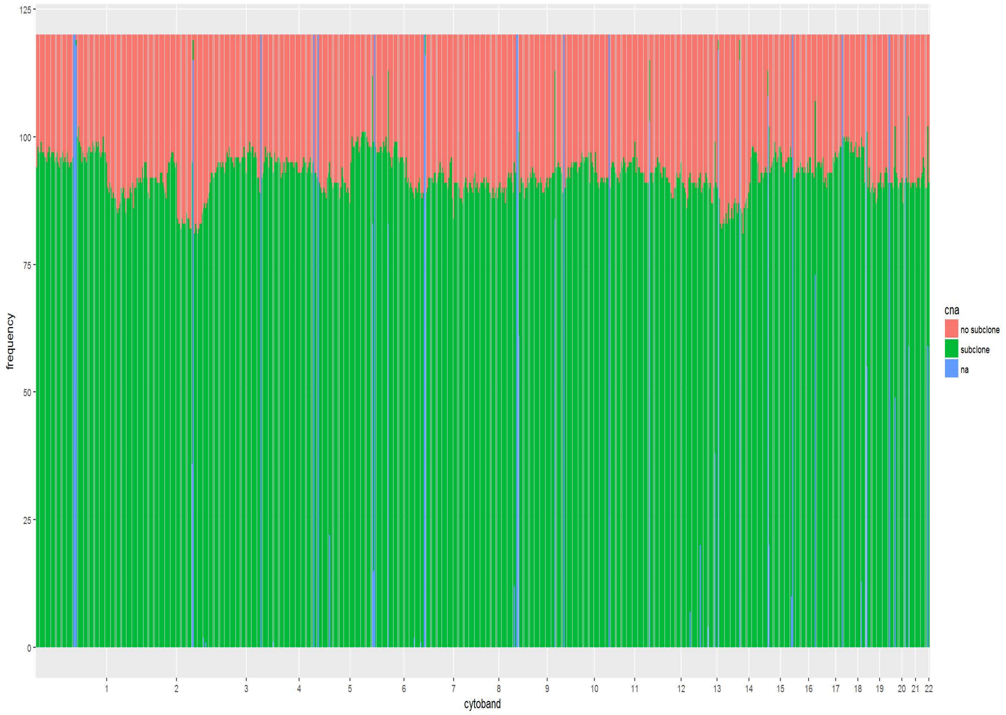


Figure D.2: Stacked histogram of subclone region profile based on subHMM for 120 TCGA renal cell carcinoma samples; na indicates no genetic information to the corresponding cytoband.s

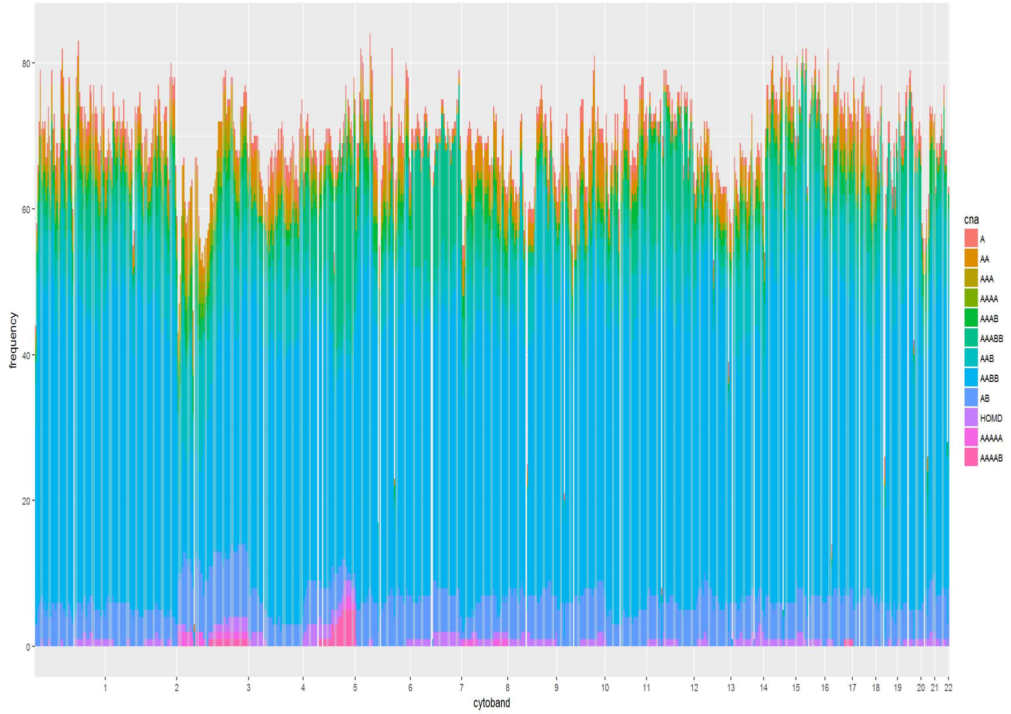


Figure D.3: Stacked histogram of subclone genotype profile based on subHMM for 120 TCGA renal cell carcinoma samples
